# In situ measurement of the isoplanatic patch for imaging through intact bone

**DOI:** 10.1101/2020.08.11.246595

**Authors:** Kayvan Forouhesh Tehrani, Nektarios Koukourakis, Jürgen Czarske, Luke J Mortensen

**Author notes:** These authors contributed equally to this work (Kayvan Forouhesh Tehrani, Nektarios Koukourakis).

## Abstract

Wavefront-shaping (WS) enables imaging through scattering tissues like bone, which is important for neuroscience and bone-regeneration research. WS corrects for the optical aberrations at a given depth and field-of-view (FOV) within the sample; the extent of the validity of which is limited to a region known as the isoplanatic patch (IP). Knowing this parameter helps to estimate the number of corrections needed for WS imaging over a given FOV. In this paper, we first present direct transmissive measurement of murine skull IP using digital optical phase conjugation (DOPC) based focusing. Second, we extend our previously reported Phase Accumulation Ray Tracing (PART) method to provide *in-situ in-silico* estimation of IP, called correlative PART (cPART). Our results show an IP range of 1-3 μm for mice within an age range of 8-14 days old and 1.00±0.25 μm in a 12-week old adult skull. Consistency between the two measurement approaches indicates that cPART can be used to approximate the IP before a WS experiment, which can be used to calculate the number of corrections required within a given field of view.

**Abstract Figure:** 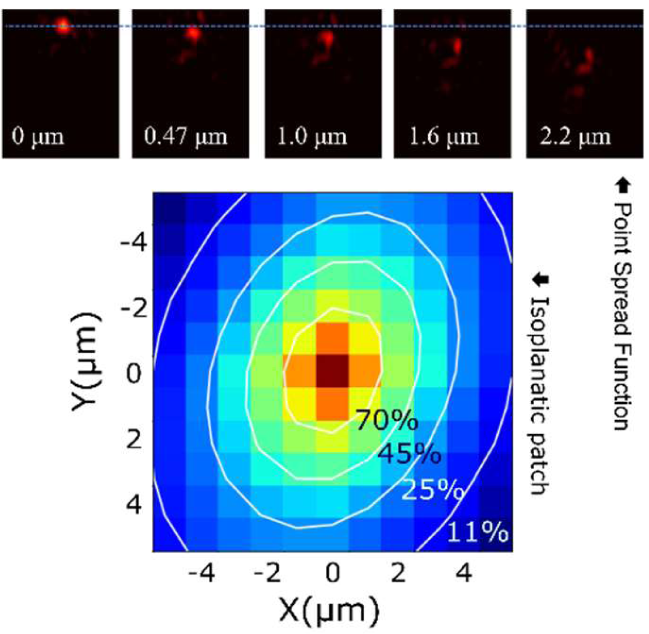

## 1. INTRODUCTION

Imaging through intact bone is highly desired for minimally-invasive monitoring of neural activity in the brain and for studying mechanisms of bone regeneration. Surgical preparations for these types of imaging frequently include an osteotomy to remove cranial bone and replace it with a transparent glass window. Although this technique is very effective, it is laborious, significantly destructive, and can alter brain or bone function due to inflammatory signaling. On the other hand, the significant optical aberrations and scattering caused by bone distort the point spread function (PSF) and make it non-trivial to image deep enough inside of a biological sample to interrogate bone or brain function [1]. Compared with bone’s mean-free-path of ∼55 μm (for 775 nm incident light), an adult (>12 weeks) murine skull is about 300 μm thick [2], while juvenile (8-14 days postnatal) skull is between 80-150 μm thick. Therefore, the layers of collagenous fibers and mineralization create a multiple-scattering [3-5] regime for light propagation even in relatively thin juvenile cranial bone, although at a smaller scale. The juvenile age range was selected because the mouse skull is still in development, and due to it being less mineralized and thinner [6] it shows a larger isoplanatic patch (IP). However since neurogenesis in cortical layers peaks at 11-17 days postnatally [7], this age group can be used in neuroimaging, as well as bone and developmental biology research.

Since multiple scattering severely limits imaging inside or through bone, several approaches have been used to extend imaging depth. One method that has been successful for the intact skull of mouse is 3-photon excitation fluorescence microscopy which exploits the longer transport-mean-free-path of near infrared light in the 1300 nm tissue transmission spectrum window [8]. Other methods that have shown success in imaging in turbid media include temporal focusing [9], speckle correlation [10, 11], *a priori* knowledge of the scattering [12-14] and artificial intelligence [15]. Skull thinning has also been successful in imaging with less distortion [16-18]. However, to achieve diffraction-limited imaging and to increase the signal to noise ratio (SNR), many of these methods could benefit from wavefront shaping (WS) methods. WS technology has previously shown that 2-photon imaging through intact skull of juvenile mouse skull is possible [19]. By engineering the wavefront and restoring the diffraction-limited PSF, WS can correct for low and high order aberrations, and enhance both the peak and width of the PSF deep inside of a sample [4, 20-30], which has many applications in the musculoskeletal system [31] and nervous system [32] imaging.

While WS improves the depth and SNR of imaging, the extent of the corrected region remains small, particularly in an anisotropically scattering tissue like bone. This effect limits the area over which a single WS correction highly correlates with its surroundings. Moving the corrected beam outside of the corrected region results in a substantial reduction of the peak and can severely distort the PSF, thereby nullifying the correction efforts. This phenomenon is known as the shift/shift scattering memory effect [33] or the IP, and is the region of shift-invariant wavefront error in a telecentric imaging system.

The IP or shift/shift memory effect can be described by the effect of physical scatterers in a turbid medium with an optical field passing through it. The scatterers cause a defined speckle pattern to the incident optical field, and if the optical field is shifted, then the speckle pattern is also shifted according to the movement of the field. In other words, there is correlation in the transmission matrices of translating fields [34], and the coherence-length of the scattering defines the extent of the area that the speckle pattern remains consistent. The Fourier conjugate of this shift/shift memory effect is known as the tilt/tilt memory effect, which was first described in waveguide geometry [35] as the angular speckle correlation effect, and described and later experimentally shown in turbid media [36-38]. Knowing the shift/shift memory effect or IP parameter is important because designing a practical WS correction experiment requires at least an estimate of the number of corrections to achieve the required field of view (FOV). Since bone is an anisotropic scattering tissue, evaluation of the tissue IP can be used to estimate the number of corrections needed for a given (FOV) using WS correction.

In this paper, we characterize the IP of *ex-vivo* juvenile (aged 8-14 days) as well as adult (>12 weeks old) murine skull experimentally using a transillumination method digital optical phase conjugation (DOPC) by focusing through the skull [39]. As we explained earlier, knowing the extent of IP becomes of the utmost importance when a correction experiment is designed. Therefore, we here introduce an extension to our previously reported Phase Accumulation Ray Tracing (PART) method to perform *in situ – in silico* estimation of IP. This method allows epi-mode estimation of wavefront phase using second harmonic generation (SHG) imaging of the bone collagen structure. In this paper we further expand our PART approach to include calculated correlation of wavefront phase within the vicinity of a corrected PSF to estimate the IP. This phase <u>c</u>orrelation-PART, which we call cPART, is capable of epi-detection mode acquisition and estimation of the IP. Therefore, it can be applied to *in vivo* experiments, which opens up new avenues for fast and efficient WS imaging through scattering tissues. We compared the results from cPART to DOPC direct measurements and found that both methods show a similar range of IP. The agreement between the results suggests that cPART can be a robust option to design the number of corrections required before WS imaging is performed *in vivo*.

## 2. METHODS

### 2.1. Digital Optical Phase Conjugation

Digital optical phase conjugation (DOPC) [40] uses direct measurement of light propagation through the scattering media, which commonly originates from a guide star (Figure 1A). Due to the limited optical access into in vivo specimen, the guide star generation for DOPC is non-trivial, which limits its applicability. However, it is an excellent method for characterization of aberration and scattering in extracted samples, like our ex-vivo skull samples, as here we have free optical access to the specimen. We created a guide star by sharply focusing light onto the specimen. The light source was a CW laser at 895 nm. The light transmitted through the skull is measured holographically and the phase conjugate is displayed on an LCoS spatial light modulator (SLM). A playback beam interacting with the SLM time reverses the light propagation direction and generates a focus through the scattering medium at the guide star position. In order to determine the extent of the IP, the correcting phase mask is shifted laterally, leading to a shift of the played back focus at the output of the sample. The correlation between the original focus and the shifted focus is computed and gives the extent of the shift. Additionally, the focus quality is analyzed using the peak to background ratio (PBR) of the playback PSF. This work defines the PBR as the ratio between the peak-intensity and the mean background in the playback PSF and can be understood as a quality measure for the DOPC procedure.

**FIGURE 1.**
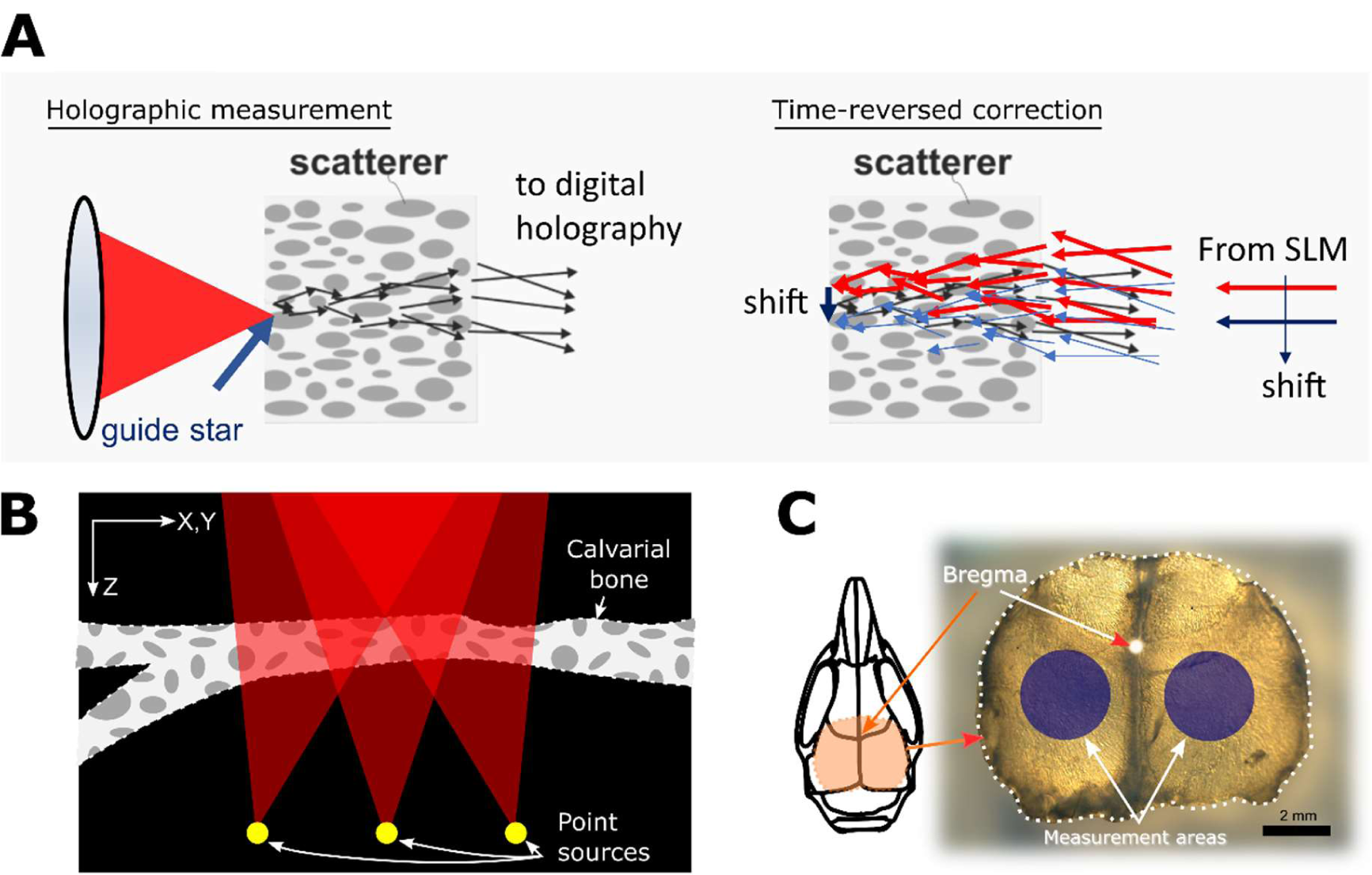
The principle of the DOPC measurement is depicted in (A). A guide star point source is created by focusing on the distal side of the skull (left). The light that transmits through the skull acts as the object beam and interferes on a digital camera with a reference beam, recording a hologram. The phase conjugate information is time-reversed using a spatial light modulator (SLM) to produce a focus at the guide star position. By shifting the correcting phase mask laterally on the SLM, the focus can be shifted; hence IP measurement can be performed. In (B) a presentation of PART method phase accumulation for each point source over a stack of SHG signal is shown. Several point sources are modeled at the bottom of the refractive-index mapped volume, and by ray tracing to the back-pupil-plane the wavefront is reconstructed. Red triangles represent the cone of focus, calculated using the numerical aperture of the objective lens. An extracted skull is shown in (C). The measurement areas are shown (blue circles) were chosen to represent the typical areas for imaging of cortex layers.

### 2.2. Correlative Phase Accumulation Ray Tracing

We previously presented PART as a method for modeling of wavefront error in tissues [1]. This method relies on an intrinsic signal generated in the tissue that can be used to calculate the approximate index of refraction of each voxel. We implement PART by acquiring second harmonic generation volumetric data of the tissue. A Ti:Sapphire source (Coherent Ultra II) tuned to 940 nm wavelength with 140 fs pulsed laser light was used for SHG imaging, and a 466/5 nm filter (Semrock) was used for light collection. Details of the microscope setup are explained elsewhere [41]. Because the SHG signal is proportional to the index of refraction of the tissue, we can use stacks of 3-dimensional images to model the tissue and hence the wavefront of the propagated light. Once the SHG image stacks were acquired, we first convert them to volumes of refraction indices. The next step is to model several point sources at the bottom of the tissue that project to the pupil plane. The angle of the cone of light propagation is determined by the numerical aperture of the microscope (Figure 1B). We acquired data in bone regions posterior to the coronal suture of the mouse skull on the left and right of the sagittal suture (Figure 1C). We used 2 skulls per age group, and in each skull 9 stacks were acquired. We used the PART analysis to output wavefronts for a grid of 11 × 11 point sources placed at the bottom of the stack. We then generated the correlation map of each wavefront *φ*_*i, j*_ with the wavefront of the center point of the 11×11 grid *φ*_*cen*_, by calculating PSF for each wavefront corrected using *φ*_*cen*_ phase. Then the Strehl ratio for each of these PSFs was calulcated using

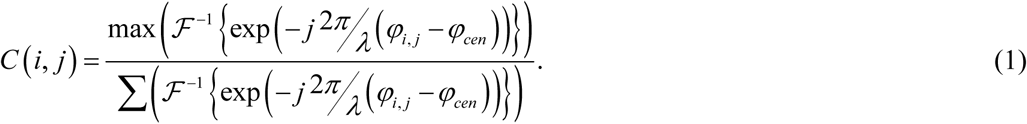

Consistent with DOPC, this method of correlation analysis produces a diffraction-limited PSF at the center (corrected using the calculated phase) and finds the extent of validity of the correcting phase mask in new locations.

### 2.3. Sample preparation

The mice aged 8-14 days postnatally (P8-P14) were euthanized according to IACUC procedures, which we defined as juvenile or pre-weaning age (< P21-P28, where P0 is the day of birth). The skulls were removed immediately and placed in 4% paraformaldehyde for fixation for 48 hours. After fixation, the skulls were washed in phosphate buffered saline (PBS), and mounted in 1% agarose gel in 35 mm Petri dishes. For measurements, the skulls were removed from the gel and washed with PBS to remove residual agarose gel. The 8-day old skulls had an average thickness of ∼80 μm, while the 14-day old skulls were >100 μm.

Although cPART can be applied to *in vivo* samples and fixation is not required, to ensure the samples maintained their integrity and remained consistent between cPART analysis and DOPC measurements, we performed fixation. Formalin-based fixation has been shown before to have minimal effect on tissue properties [42]. We measured the scattering mean-free-paths of a tissue before and after correction (Figure 2), and found 39.4 ± 2.4 µm and 40.6 ± 2.2 µm respectively, using 775nm light. We made an effort to image similar areas for these measurements, and we find a 6.8% difference between the two measurements, which shows an insignificant difference.

**FIGURE 2.**
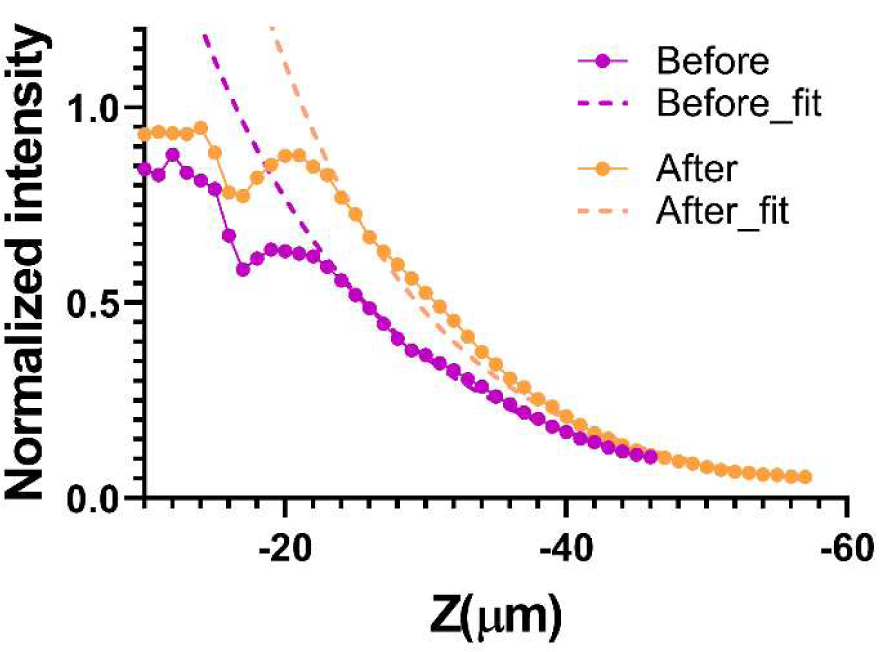
Comparison of the SHG signal decay before and after fixation. The mean-free-paths were 39.4 ± 2.4 µm and 40.6 ± 2.2 µm, using 775nm light, before and after fixation, respectively.

For *in vivo* imaging, the mice were anesthetized using isoflurane (initially 4% and 1.9% during imaging, with 100 ml/min oxygen flow rate), and restrained using a 3D printed stereotaxic holder. One hour before surgery a 1mg/kg dose of Meloxicam was injected subcutaneously to the mouse, and five minutes before making an incision a 5mg/kg dose of bupivacaine was locally applied as analgesia. A V-shaped incision was made on the scalp from between the eyes toward both the ears to create a flap. The periosteum layer was removed, and the area of imaging was cleaned using a cotton swab; immediately sterile phosphate buffered saline (PBS) was applied to the incision site. The animal was placed under the microscope objective, and sterile PBS was added to fill the gap between the skull and the objective lens. After the imaging session, the animal was euthanized using CO2 and cervical dislocation. All animal procedures and experiments were approved by the UGA Institutional Animal Care and Use Committee (IACUC).

## 3. RESULTS AND DISCUSSION

### 3.1. DOPC direct measurement

We performed the DOPC procedure as described in section 2.1 and obtained a focus after DOPC through the skull (Figure 3A, left). The correcting phase mask was shifted laterally leading to a shift of the focus, and at each point we measured the peak to mean background ratio (PBR) of the PSF. Since the quality of the PSF depends on how much of the light is concentrated in the PSF relative to the total energy, the ratio of the desired intensity (peak intensity) and undesired speckles (mean background) produces a useful metric for this analysis. As expected, increasing the shift distance from the corrected spot leads to a reduction in PBR and deterioration of the PSF quality, meaning that more energy is spread in the speckle background rather than the main lobe of the PSF. For each sample this measurement was repeated at 6-10 lateral positions.

**FIGURE 3.**
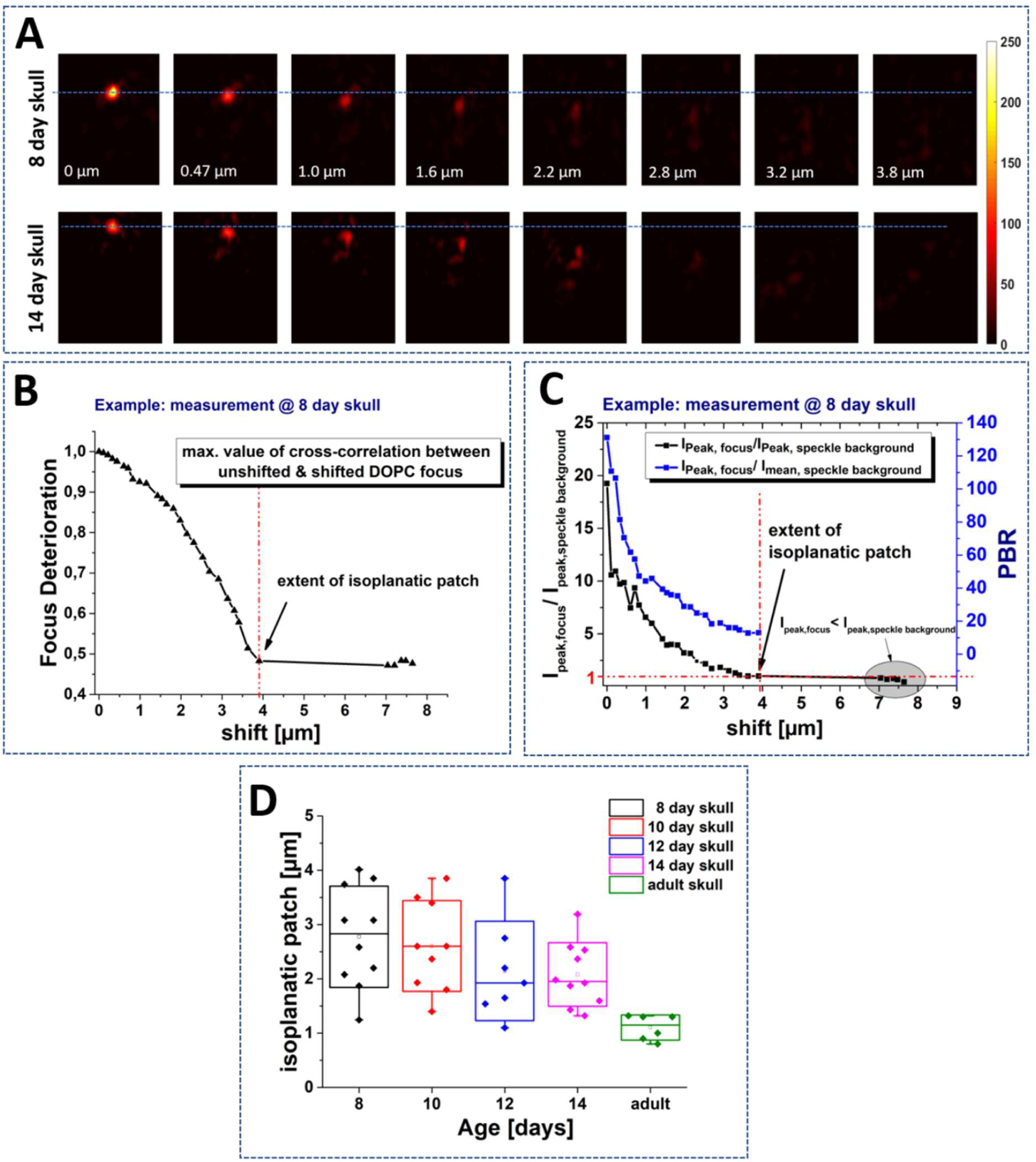
DOPC measurement of bone IP. A) After DOPC time-reversed focus is created through the skull (position 0 µm). The phase mask is shifted laterally resulting in a shift of the focus, which shows a deteriorating quality with the shift. Exemplary results are shown for two cases of 8-day and 14-day skull. The behavior for both ages is comparable. B) The correlation between the unshifted focus and the shifted light-distribution shows a decorrelation behavior. We count the “point of decorrelation” and the extent of the memory effect as the shifting length at which the peak intensity in the focus is higher than the maximum intensity in the speckle background C). The shift length of the decorrelation point is plotted in D) for different ages and at different positions.

To quantify the extent of the IP, we use the normalized cross-correlation between the focus at zero shift and the shifted focus. For each PSF shift-distance, we obtain a cross-correlation measurement, which describes the persistency of the focus at that point (Figure 3B). To determine the extent of the IP, we find the point where the peak of the background speckle intensity has a higher intensity than the expected PSF position (Figure 3C gray area). This means that the maximum of the cross-correlation is not at the expected shifted PSF position, but is distributed randomly across the field and remains nearly constant upon further shift. The exact value of the correlation was calculated as focus deterioration (Fig 3C), and depends on the speckle-background amplitude and initial quality of the focus and is not expected to drop to zero.

Two skulls per age were measured at different positions. The resulting IP measurements are plotted in Figure 3D. Although there appeared to be a tendency towards smaller IP size as the mice matured (P8:2.7 µm ± 1.3 µm, P14: 2.0 µm ± 1.15 µm), and a reduction in the variance for adult mice, the effect of age on IP measurements did not show significant difference among the juvenile skulls (1-way ANOVA P-value=0.2). The resulting overall range of IP is about 1.1 to 4 μm.

For comparison, experiments were also conducted on adult skulls, leading to a reduced extent of the IP range of 1.1 μm ± 0.2 μm. This comes with a higher thickness of the skull, which is approximately >350 µm. However, it is not distinguishable how much of this reduction is attributed to the higher thickness versus the contribution of other structural changes in the bone (increased mineral density, bone marrow cavity alterations, changes in vascularity, etc.). It is also essential to think about the quality of the PSF within an IP. Although we calculate the extent of IP by finding the maximum range of correlation, the PBR degrades towards the edge of the IP (Figure 3C), and so PSFs at the end of this range have low PBR values and might not be usable for imaging implementations. Therefore, we create an additional measure to take into account the ratio between the maximum intensity in the focus and the maximum intensity occurring in the speckle-background. We assume the usable range to be up to the point where *I* _*peak. focus*_ = 3 × *I* _*peak. speckle* − *background*_ (at a PBR of approximately 30). In the presented example of a skull of P8, we find a usable shift-range of about 2.2 µm. The exact values may vary as they depend on several experimental and sample parameters. Therefore, although these values may not be generally applicable, they provide an approximate range for the skulls of this age range.

### 3.2. cPART modeling

We used PART simulations to produce wavefronts over a grid of equally spaced 11×11 point-sources in a single image plane with a lateral interval of 0.9 μm. Representative examples of 8-day old and 14-day old wavefront unwrapped phases show significant impact of the tissue on the wavefront (Figure 4A). After calculating the correlations, we applied a 2D Gaussian fit with rotating axis *θ*:

**FIGURE 4.**
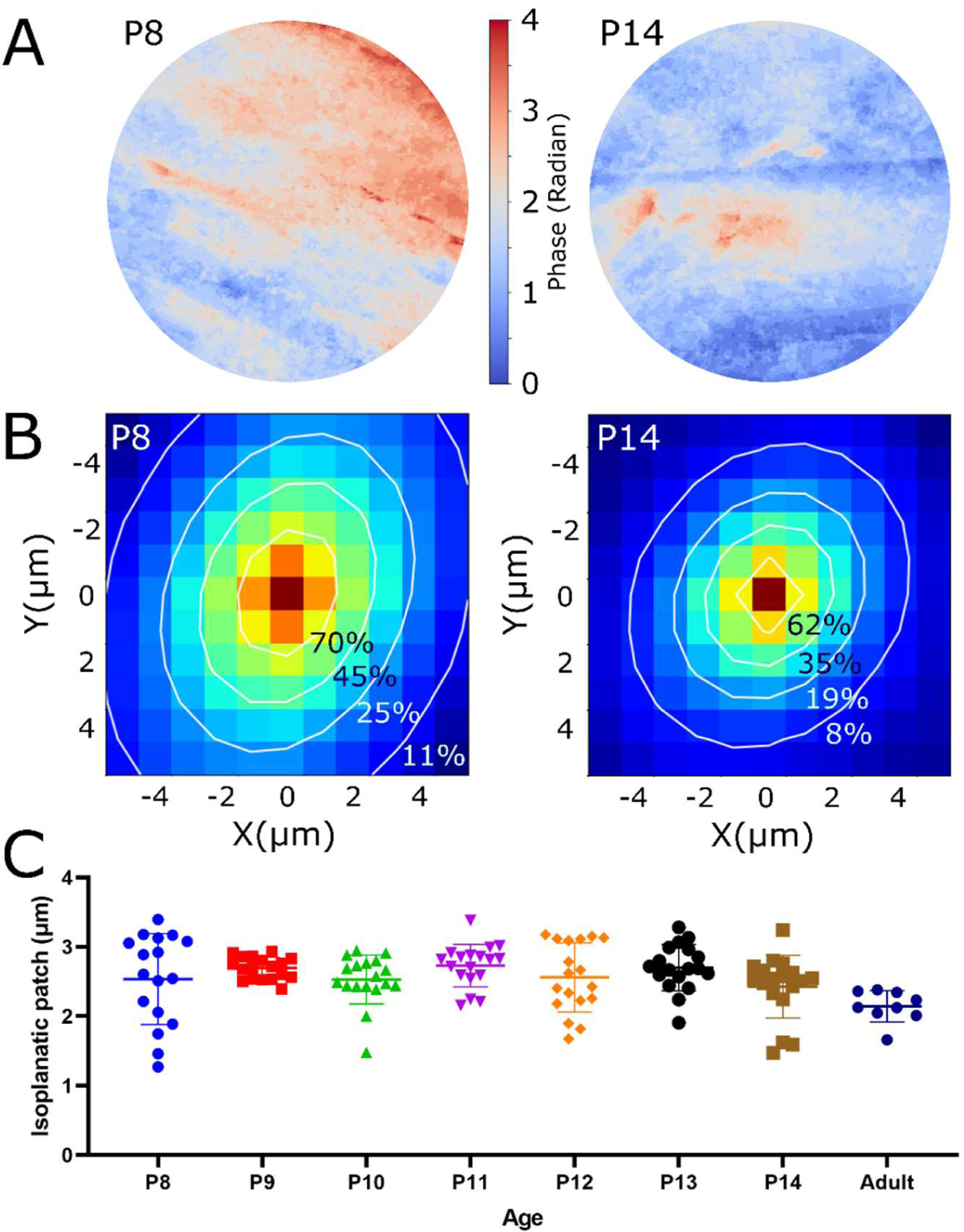
PART simulation of bone wavefront error and the isoplanatic patch. (A) Examples of PART simulation of wavefront phase at the back pupil (without phase wrapping) for 8 and 14 day old mouse skulls. The extent of the pupils are determined using the numerical aperture of the objective lens and the wavelength of light. (B) Correlation of simulated pupils from multiple originating points and Gaussian fit (white contour lines) over the correlation patterns. (C) Using the Gaussian fitting the FWHM of the correlation is calculated and shown for each age group.

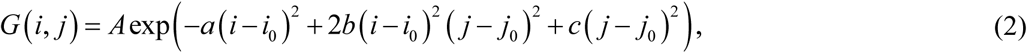

where 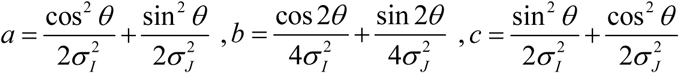, and *A* is the amplitude.

The SHG stacks used for this study had a thickness range of between 80-150 µm. Correlation maps and contour lines of the applied fittings show the difference between the size of a representative correlation in the P8 animal versus P14 (Figure 4B). Using the sigma values of the major and minor axes of the Gaussian fitting we calculated the extent of IP by calculating the full width and half maximum (FWHM) of their hypotenuse and compare the extent of the IPs of skulls of different ages (Figure 4C). These results show a range of 2.53 μm ±0.6 patch for 8-day old, to 2.42 μm ±0.4 for 14-day old skulls. The absolute range between all age groups is 1.27 μm to 3.39 μm. Confirmed by one-way ANOVA test, the mean values of the P8-P14 measurement groups do not show a significant difference (2.59 μm ±0.1, P-value = 0.258); however, their standard deviations are significantly different (P-value < 0.001), which is due to the range being reduced as the mice age and have thicker skulls. This variation indicates that different structures that occur throughout regions of the bone, such as vasculature, bone marrow cavities, and topology of the skull, start to become more consistent as the mice age. Since our cPART results are consistent with our direct DOPC measurements, it suggests that cPART can be used to estimate the extent of IP and by extension, the number of corrections needed over a given FOV. We further performed IP measurement on *in vivo* adult skull of a 12 week old mouse (Figure 4C), which shows a range of 2.14 μm ±0.2 (the adult sample presented here is different from the DOPC adult skull measurement).

A limitation of cPART is that SHG imaging’s maximum achievable depth is limited by the transport-mean-free-path of the tissue (inversely proportional to scattering coefficient and 1-g tissue coefficient parameter) and thus, this technique is not applicable for full depth estimation in thicker tissues. Nevertheless, for samples of appropriate thickness or desired imaging depths (such as the juvenile murine skulls) the use of cPART enables the epi-detected approximation of the IP, which has not previously been reported. This capability could facilitate accurate measurement of the IP size during a relatively large FOV wavefront shaping imaging deep inside of bone and brain. In addition to WS imaging, our findings suggest that by characterizing the bone wavefront distortions using cPART *in situ*, it could be possible to assess both collagen organization and bone density quality metrics simultaneously. This could be used to understand severity and treatments efficacy for diseases that affect mineralization as well as the collagenous structure of the bone, such as hypophosphatasia, osteogenesis imperfecta, x-linked hypophosphatemia, or osteoporosis.

## 4. CONCLUSION

In conclusion, this work presents measurement of the IP and a simulation method for its estimation in mouse skulls of 8-14 day old. This age range was selected because they have relatively thinner skulls than adult mouse, which is a suitable choice for imaging inside or through their skull. This age range is particularly interesting for imaging, because the neurogenesis in the cortical layers peak in P12-P17[7], but bone maturation has not peaked yet[6], which makes it suitable for neuroimaging, as well as bone and developmental research. We performed DOPC measurements to directly measure the IP of the bone. Because the DOPC method requires transillumination of the sample it is not applicable to in-vivo determination of the IP, so we extended our previous simulation method to allow phase correlation PART (cPART) and estimate the IP in intact samples. Both techniques show in good agreement that an IP range of about 1-4 μm is expected, therefore if a FOV of 20 × 20 μm^2^ is desired to image in the best case 25 corrections must be made, and in the worst case 400 corrections. This range can be estimated using cPART modeling prior to wavefront correction. Because this method works in epi-illumination mode, it can be applied to *in vivo* imaging of mouse neural activity or brain imaging through intact or thinned skull. We showed that the mean value of the measurements between the age groups are mostly consistent, however, the 8-day old mice are more likely to have a larger memory effect which significantly reduces the number of corrections required for a larger FOV. This method of analysis can also be used for the characterization of bone, which is likely reflective of the bone density and structure. In future we will extend the presented analysis to mice with congenital diseases such as hypophosphatasia and osteogenesis imperfecta to assess bone density and structure.

## ACKNOWLEDGMENT

LM and KF were supported in part by the National Science Foundation (1706916) and the National Institutes of Health (R21EB027802); and the work was supported by grants to LM from the Center for Regenerative Engineering and Medicine (REM) and the Soft Bones Foundation. JC and NK were supported by Reinhart Koselleck Project of German Science Foundation (CZ55-30).

## Abbreviations

DOPC: Digital Optical Phase Conjugation
FOV: Field of View
IP: Isoplanatic Patch
PART: Phase Accumulation Ray Tracing
PSF: Point Spread Function
SNR: Signal to Noise Ratio
WS: Wavefront Shaping

## REFERENCES

1. Tehrani, K.F., P. Kner, and L.J. Mortensen, Characterization of wavefront errors in mouse cranial bone using second-harmonic generation. Journal of Biomedical Optics, 2017. 22(3): p. 036012–036012.

2. Soleimanzad, H., H. Gurden, and F. Pain, Optical properties of mice skull bone in the 455-to 705-nm range. Journal of Biomedical Optics, 2017. 22(1): p. 010503.

3. Berkovits, R. and S. Feng, Correlations in coherent multiple scattering. Physics Reports, 1994. 238(3): p. 135–172.

4. Vellekoop, I.M. and A.P. Mosk, Phase control algorithms for focusing light through turbid media. Optics Communications, 2008. 281(11): p. 3071–3080.

5. Jarry, G., F. Henry, and R. Kaiser, Anisotropy and multiple scattering in thick mammalian tissues. Journal of the Optical Society of America A, 2000. 17(1): p. 149–153.

6. Wei, X., et al., Postnatal Craniofacial Skeletal Development of Female C57BL/6NCrl Mice. Front Physiol, 2017. 8: p. 697.

7. Finlay, B.L. and R.B. Darlington, Linked regularities in the development and evolution of mammalian brains. Science, 1995. 268(5217): p. 1578–84.

8. Wang, T., et al., Three-photon imaging of mouse brain structure and function through the intact skull. Nature Methods, 2018.

9. Escobet-Montalbán, A., et al., Wide-field multiphoton imaging through scattering media without correction. Science Advances, 2018. 4(10): p. eaau1338.

10. Yang, X., Y. Pu, and D. Psaltis, Imaging blood cells through scattering biological tissue using speckle scanning microscopy. Optics Express, 2014. 22(3): p. 3405–3413.

11. Hofer, M., et al., Wide field fluorescence epi-microscopy behind a scattering medium enabled by speckle correlations. Optics Express, 2018. 26(8): p. 9866–9881.

12. Guo, K., et al., 13-fold resolution gain through turbid layer via translated unknown speckle illumination. Biomedical Optics Express, 2018. 9(1): p. 260–275.

13. Chang, J. and G. Wetzstein, Single-shot speckle correlation fluorescence microscopy in thick scattering tissue with image reconstruction priors. 2018. 11(3): p. e201700224.

14. Guo, C., et al., Imaging through scattering layers exceeding memory effect range by exploiting prior information. Optics Communications, 2019. 434: p. 203–208.

15. Cheng, S., et al., Artificial intelligence-assisted light control and computational imaging through scattering media. 2019. 12(04): p. 1930006.

16. Li, Y., U. Baran, and R.K. Wang, Application of Thinned-Skull Cranial Window to Mouse Cerebral Blood Flow Imaging Using Optical Microangiography. PLOS ONE, 2014. 9(11): p. e113658.

17. Dorand, R.D., et al., Comparison of intravital thinned skull and cranial window approaches to study CNS immunobiology in the mouse cortex. IntraVital, 2014. 3(2): p. e29728.

18. Turcotte, R., et al., Characterization of multiphoton microscopy in the bone marrow following intravital laser osteotomy. Biomedical Optics Express, 2014. 5(10): p. 3578–3588.

19. Park, J.-H., W. Sun, and M. Cui, High-resolution in vivo imaging of mouse brain through the intact skull. Proceedings of the National Academy of Sciences, 2015. 112(30): p. 9236–9241.

20. Tao, X., et al., Adaptive optical two-photon microscopy using autofluorescent guide stars. Optics Letters, 2013. 38(23): p. 5075–5078.

21. Zhang, X. and P. Kner, Binary wavefront optimization using a genetic algorithm. Journal of Optics, 2014. 16(12): p. 125704.

22. Vellekoop, I.M., Feedback-based wavefront shaping. Optics Express, 2015. 23(9): p. 12189–12206.

23. Kong, L. and M. Cui, In Vivo Deep Tissue Imaging via Iterative Multiphoton Adaptive Compensation Technique. Ieee Journal of Selected Topics in Quantum Electronics, 2016. 22(4).

24. Li, Q., et al., Woofer-tweeter adaptive optical structured illumination microscopy. Photonics Research, 2017. 5(4): p. 329–334.

25. Tao, X., et al. A three-photon microscope with adaptive optics for deep-tissue in vivo structural and functional brain imaging. 2017.

26. Tehrani, K.F., et al., Adaptive optics stochastic optical reconstruction microscopy (AOSTORM) using a genetic algorithm. Optics Express, 2015. 23(10): p. 13677–13692.

27. Tehrani, K.F., et al., Adaptive optics stochastic optical reconstruction microscopy (AOSTORM) by particle swarm optimization. Biomedical Optics Express, 2017. 8(11): p. 5087–5097.

28. Czarske, J.W., et al., Transmission of independent signals through a multimode fiber using digital optical phase conjugation. Optics Express, 2016. 24(13): p. 15128–15136.

29. Koukourakis, N., et al., Wavefront shaping for imaging-based flow velocity measurements through distortions using a Fresnel guide star. Optics Express, 2016. 24(19): p. 22074–22087.

30. Kuschmierz, R., et al., Self-calibration of lensless holographic endoscope using programmable guide stars. Optics Letters, 2018. 43(12): p. 2997–3000.

31. Tehrani, K.F., et al., Two-photon deep-tissue spatially resolved mitochondrial imaging using membrane potential fluorescence fluctuations. Biomedical Optics Express, 2018. 9(1): p. 254–259.

32. Schlüßler, R., et al., Mechanical Mapping of Spinal Cord Growth and Repair in Living Zebrafish Larvae by Brillouin Imaging. Biophysical Journal, 2018. 115(5): p. 911–923.

33. Osnabrugge, G., et al., Generalized optical memory effect. Optica, 2017. 4(8): p. 886–892.

34. Judkewitz, B., et al., Translation correlations in anisotropically scattering media. Nature Physics, 2015. 11(8): p. 684–689.

35. Feng, S., et al., Correlations and Fluctuations of Coherent Wave Transmission through Disordered Media. Physical Review Letters, 1988. 61(7): p. 834–837.

36. Freund, I., M. Rosenbluh, and S. Feng, Memory Effects in Propagation of Optical Waves through Disordered Media. Physical Review Letters, 1988. 61(20): p. 2328–2331.

37. Berkovits, R., M. Kaveh, and S. Feng, Memory effect of waves in disordered systems: A real-space approach. Physical Review B, 1989. 40(1): p. 737–740.

38. Berkovits, R. and M. Kaveh, Time-reversed memory effects. Physical Review B, 1990. 41(4): p. 2635–2638.

39. Koukourakis, N., M. Kreysing, and J. Czarske. Wavefront shaping method to focus through mouse skull. in Imaging and Applied Optics 2018 (3D, AO, AIO, COSI, DH, IS, LACSEA. LS&C, MATH, pcAOP). 2018. Orlando, Florida: Optical Society of America.

40. Vellekoop, I.M., M. Cui, and C. Yang, Digital optical phase conjugation of fluorescence in turbid tissue. Applied Physics Letters, 2012. 101(8): p. 081108.

41. Tehrani, K.F., et al., Five-dimensional two-photon volumetric microscopy of in-vivo dynamic activities using liquid lens remote focusing. Biomedical Optics Express, 2019. 10(7): p. 3591–3604.

42. Roberta, G., et al., Effects of tissue fixation on coherent anti-Stokes Raman scattering images of brain. Journal of Biomedical Optics, 2013. 19(7): p. 1–8.

